# *magpie*: a power evaluation method for differential RNA methylation analysis in N6-methyladenosine sequencing

**DOI:** 10.1101/2023.09.08.556896

**Authors:** Zhenxing Guo, Daoyu Duan, Wen Tang, Julia Zhu, William S. Bush, Liangliang Zhang, Xiaofeng Zhu, Fulai Jin, Hao Feng

## Abstract

**Motivation:** Recently, novel biotechnologies to quantify RNA modifications became an increasingly popular choice for researchers who study epitranscriptome. When studying RNA methylations such as N6-methyladenosine (m^6^A), researchers need to make several decisions in its experimental design, especially the sample size and a proper statistical power. Due to the complexity and high-throughput nature of m^6^A sequencing measurements, methods for power calculation and study design are still currently unavailable.

**Results:** We propose a statistical power assessment tool, *magpie*, for power calculation and experimental design for epitranscriptome studies using m^6^A sequencing data. Our simulation-based power assessment tool will borrow information from real pilot data, and inspect various influential factors including sample size, sequencing depth, effect size, and basal expression ranges. We integrate two modules in *magpie*: (i) a flexible and realistic simulator module to synthesize m^6^A sequencing data based on real data; and (ii) a power assessment module to examine a set of comprehensive evaluation metrics.

**Availability:** The proposed power assessment tool *magpie* is publicly available as a R/Bioconductor package at: https://bioconductor.org/packages/magpie/.

## Introduction

RNA methylation represents another layer of epigenetic regulation in addition to the well-studied DNA methylation and histone modification. Among different types of RNA methylation, N6-methyladenosine, i.e. m^6^A, is the most common form. It has been identified as one of the post-transcriptional regulatory markers on mRNA, rRNA, tRNA, circRNA, miRNA and longnoncoding RNA, and plays important roles in regulating pre-RNA splicing, RNA translation, stability, and degradation (Wang et al., 2014; Geula et al., 2015; Oerum et al., 2021). The effects of m^6^A suggest its involvement in multiple cellular processes such as cell differentiation and reprogramming (Lasman et al., 2020; Chen et al., 2015). Studies also suggest linkages between the dysregulation of m^6^A and many human diseases such as cancers and neural disorders (Geula et al., 2015; Chen et al., 2019; Lan et al., 2019).

MeRIP-seq/m^6^A-seq was developed to characterize transcriptome-wide m^6^A profiles (Dominissini et al., 2012; Meyer et al., 2012). This technique typically relies on immunoprecipitation of m^6^A-containing RNA fragments (m^6^A-IP), followed by high-throughput next generation sequencing. These samples are generally referred to as the IP (immunoprecipitated) samples. In addition to IP samples, cDNA libraries are also prepared for input control mRNAs to measure the background mRNA abundance. These input controls are essentially the transcriptome from regular RNA-seq. The m^6^A methylation level, for each region, is then quantified by the enrichment of IP over input, roughly the normalized ratio between IP and input control counts. If the m^6^A enrichment is significantly high, then the called peak of that region suggests an underlying m^6^A residue. MeRIP-seq is becoming a popular and indispensable tool to profile transcriptome-wide m^6^A, since the invention of this technique.

One important characteristic of MeRIP-seq is its large technical variability among different samples, partially due to the independent immunoprecipitation procedure. Such technical artifacts lead to erroneous peak calling of methylated regions. This problem becomes prominent in studies with small sample sizes (McIntyre et al., 2020), which is often the case given the high expenses associated with the current experimental protocols. In m^6^A-seq2 (Dierks et al., 2021), a single IP experiment is performed on the pooled RNAs of all samples, where RNAs from different samples are uniquely barcoded and demultiplexed after sequencing. The multiplexed profiling procedures could reduce the deviations; however, technical variability is still an unignorable component in m^6^A data analysis.

To study the biological implications of m^6^A, one fundamental task is to identify the Differentially Methylated Regions (DMRs) across different conditions. Although several DMR detection methods have been developed (Tang et al., 2021; Zhang et al., 2019; Guo et al., 2022) and evaluated (Duan et al., 2023) in either MeRIP-seq or m^6^A-seq2 experiments, the sample size calculation with their associated statistical power remains an open question due to the complexities of sequencing experiments. Further, due to the uniqueness in data structure, power analysis tools developed for other types of sequencing data cannot be applied to MeRIP-seq and m^6^A-seq2 experiments. To be specific, the input control data from MeRIP-seq and m^6^A-seq2 are from RNA-seq experiments, where there is a mean-variance dependence. The effect size is hidden in the ratio between IP and input counts, not the input data alone. In addition, because the count coverage of each gene depends on both its expression level and sequencing depth of the whole sample, the sequencing depth is another experimental factor that affects the variance estimate and thus the accuracy of a site to be detected as a significant DMR; therefore, it needs to be considered in power analysis. Currently, with the increasing popularity of epitranscriptome studies, an appropriate statistical procedure is needed to optimize the sample size and assist researchers with proper power analysis. To our best knowledge, no method is currently available.

Here, we propose a comprehensive power evaluation method named ***magpie*** (**m**^6^**A g**enome-wide differential analysis **p**ower **i**nferenc**e**). ***magpie*** first learns characteristics of real data, and then synthesizes data that mimics the real data well. In simulations, ***magpie*** allows for the adjustment of sample size, sequencing depth and effect size. It can evaluate the epitranscriptome study design using multiple metrics including sensitivity, specificity, precision, false discovery rate, and more. Our proposed method is the first available tool to guide the practical experimental design by comprehensively investigating the relationship between statistical metrics and associated factors in m^6^A differential analysis. ***magpie*** is publicly available as an R/Bioconductor package at https://bioconductor.org/packages/magpie/.

## Materials and Methods

### An Overview of *magpie*

We assess the effect of experimental design on the power of DMR detection purely based on simulations, where the whole procedure is divided into two components. First, ***magpie*** preprocesses .*bam* files from MeRIP-seq sequencing and obtains read counts in candidate regions from all samples (Fig. 1), where candidates are identified with conditional binomial tests. With the counts from the identified candidate regions, ***magpie*** simulates count matrices for both IP and Input samples with a Gamma-Poisson model. Parameters involved are estimated from the candidates to mimic the actual MeRIP-seq data in aspects of marginal distribution read counts, and the distribution of biological dispersion in methylation levels (Fig. 1). With data simulated, we then evaluate power and error rates on them (Fig. 1). The two components, Gamma-Poisson simulation and power assessment, are independent so that ***magpie*** allows the assessment on data by different simulation strategies.

**Figure 1.**
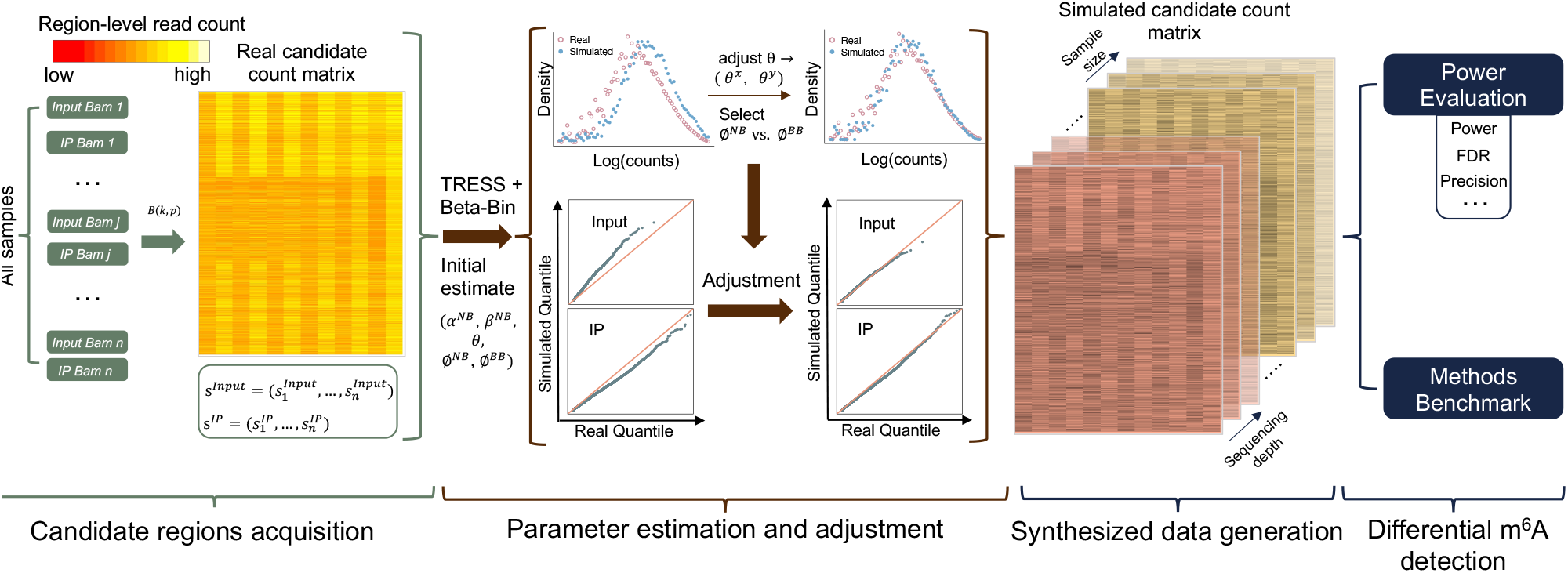
Overview of ***magpie. magpie*** provides power evaluation for differential m^6^A methylation analysis. It takes pilot MeRIP-seq data as the input. Based on the pilot data, it obtains candidate regions, estimates key parameters, and conducts real-data-based simulations for statistical power evaluation.

### Data Generative Model

Here we describe how ***magpie*** simulates the MeRIPseq count matrices given existing real MeRIP-seq data from different conditions. ***magpie*** processes .*bam* files by splitting the transcriptome into bins, aggregating read counts, and testing for significance of IP enrichment over Input. Using a bump-finding algorithm, significant bins are combined into candidate regions. ***magpie*** then focus on these candidates in simulations, as other regions lack IP enrichment and biological relevance. Suppose there are in total *N* pairs of IP and Input samples from all conditions, and *M* candidate DMRs generated after preprocessing. Let *X*_*ij*_ and *Y*_*ij*_ denote input and IP counts in candidate DMR *i* from sample *j*. We assume that 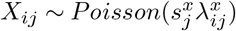 and 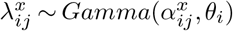 Similarly, 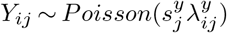 and 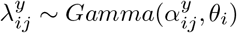. Here 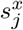 and 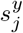 represent the normalizing factors for input and IP samples, such as the library sizes. 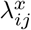 and 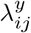 are normalized poisson rates. 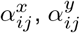, and *θ*_*i*_ are the shape and scale parameters of corresponding gamma distributions. Given above assumptions, naturally 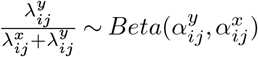.

Further, denote 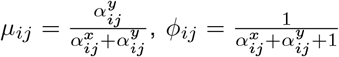, then marginally,

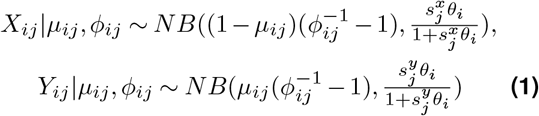

In above equations, *μ*_*ij*_ and *φ*_*ij*_ represent the mean and dispersion of the methylation level for candidate region *i* in sample *j*.

We begin by simulating size factors, for which we directly use the values estimated from real data:

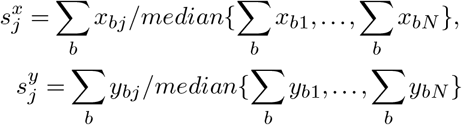

where *x*_*bj*_ and *y*_*bj*_ are read counts in bin *b* from the *j*th Input and IP samples.

Next, for each candidate region *i*, ***magpie*** simulates a baseline methylation level *μ*_*i*_ or equivalently 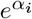 in the structure of 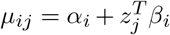 where *z*_*j*_ contains the attributes of sample *j*, and *β*_*i*_ represents corresponding effects. To do that, we randomly sample *α*_*i*_ from one parametric distribution, or from its empirical distribution estimated from real data.

After simulating the baseline methylation, we simulate 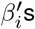 s for all regions. Because we can hardly know the actual number of DMRs and their degree of differential methylation, specific settings are adopted based on both reasonable assumptions and empirical observations. First, ***magpie*** sets the proportion of DMRs as 10%, assuming that DM is present in only a small subset of regions in most experiments. Then, for non-DMRs, *β*_*i*_ = 0. For DMR 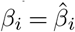 if its estimated effect size 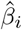 is greater than the 50% quantile of all regions. Otherwise, *β*_*i*_ ∼ *U* (1, 2). The exact quantile and the range of uniform distribution can be adjusted by users to explore a broad range of effect sizes.

For dispersion *φ*_*ij*_, we can simulate it again from a parametric distribution or sample from empirical distributions. To ensure the robustness, the empirical distribution can be estimated by TRESS from raw counts or by Beta-binomial regressions from normalized counts. Denote 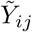 and 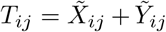 as the normalized IP and total counts, the Beta-Binomial regressions are established as follows:

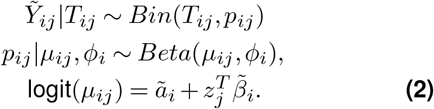

where *μ*_*ij*_ and *φ*_*i*_ represent the mean and dispersion of methylation level. As noted, for the convenience of estimation, above Beta-binomial regressions (as well as TRESS) assume *φ*_*ij*_ = *φ*_*i*_ for all *j*.

Empirically for the same real dataset, 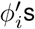 estimated by the Negative-binomial model in TRESS are usually greater than those estimated by Beta-binomial regressions. Without golden truth, our setting for *φ*_*i*_ relies on a data driven approach. Specifically, by comparing to the real data, ***magpie*** will calculate a KL-divergence for each of the synthetic counts by model in Eq. (1). Those 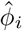 resulting in a significantly lower KL divergence between simulated data and real data will be kept for final data generation. If there is no significant difference in KL-divergence, 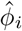 estimated by NB will be adopted.

Lastly, we simulate the scale parameter *θ*_*i*_ in Eq. (1). Again, it can be simulated by parametric distributions, or sampled directly from empirical distributions. For the parametric distribution, we set its mean as a function of *φ*_*i*_ observed in the previous peak detection method (Guo et al., 2022). No matter the strategy employed, the first-round generated 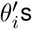 will be further scaled by the fold change between real and first-round simulated counts. Such an adjustment again helps to reduce the disparity between the simulated and real distributions, thereby improving the reliability of the results in followup power assessments.

### DMR Detection

After generating the simulated read counts in candidate DMRs, an existing software developed for MeRIP-seq is applied to detect DMRs. We implement an interface for calling TRESS and exomePeak2 (Tang et al., 2021). Users can define other DMR detection methods and plug into the procedure. Each method reports test statistics, P-values and FDRs for all candidate regions. These results are then used for the downstream power assessment.

### Power Assessment Measures

We adopted several evaluation metrics in the statistical power assessment for differential analysis using MeRIP-seq data. These metrics include classic criteria in hypothesis testings such as the false discovery rate (FDR), power, and precision. We also inspected the false discovery cost (FDC, defined below) and targeted power (Wu et al., 2015), aiming to provide a comprehensive statistical power evaluation.

Because not all DMRs are of biological interest to us, especially those with low effect sizes, we introduce a cutoff ∆ for the effect size *β*. Only those DMRs with |*β*| ≥ ∆ are considered as ‘targeted DMRs’, which are of biological interest in research. We denote the number of non-DMRs, non-targeted DMRs, and targeted DMRs as *R*_0_, *R*_1_, and *R*_2_, respectively. Suppose *T*_*r*_ represents the testing result of region *r*, where *T*_*r*_ = 1 denotes the discovered DMR, and *T*_*r*_ = 0 otherwise. The confusion matrix in the DMR detection is summarized in Table 1.

**Table 1.**
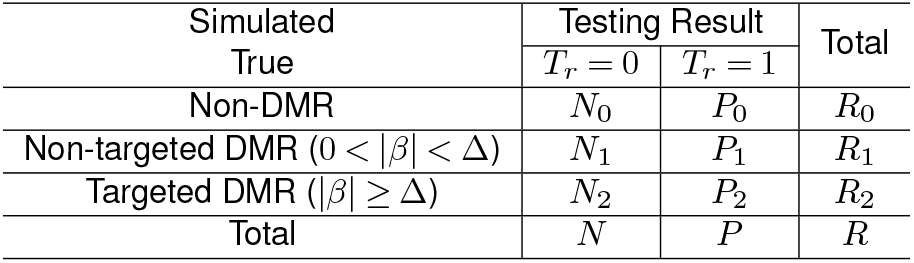
The confusion matrix in m^6^A DMR detection, when taking biological significance into consideration.

The false discovery rate (FDR) and precision are statistical metrics that jointly provide insights into the balance between true and false discoveries among the significant features. In practical experiments, such as MeRIPseq, researchers often focus on the top detected DMRs by their chosen methods for further analyses. As such, the proportion of true DMRs among the significants is essential to ensure the reliability of their downstream analyses. Thus, type I error control is often a prime task. In this context, FDR and precision are defined as 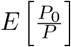 and 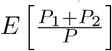, respectively. Statistical power is defined, naturally, as 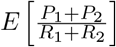 To investigate the power of detecting targeted DMRs that are biologically interesting with |*β*| ≥ ∆, targeted power is introduced and defined as 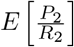 . To better illustrate the tradeoff between false positives and true positives, we propose an additional metric, False Discovery Cost (FDC),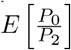, which is defined as the expected number of false positives per targeted true positive. The rationale behind this is straightforward: this cost is the expected number of false discoveries, per true discovery we are interested in.

Finally, our proposed evaluation framework allows for the examination of aforementioned metrics using simulations under various combinations of sample size, sequencing depth, input expression stratum, and FDR threshold. Each user-defined scenario is repeated for 100 times, and these metrics are computed and averaged to generate empirical estimations.

### Implementation

Given a MeRIP-seq dataset in .*bam* files, various experimental scenarios (such as sample size, sequencing depth, FDR threshold, etc.), and a chosen differential methylation testing method, ***magpie*** generates evaluation results for each proposed study design. Functions incorporated in ***magpie*** allow users to export these results in an .*xlsx* file, and to visualize them through line plots. Users have the option to provide small pilot data, which could include only several chromosomes. We would estimate major parameters from these pilot data, to guide larger-scale simulations for power evaluation for future experimental designs. Alternatively, when pilot MeRIP-seq datasets are unavailable or unattainable, the function *quickPower* can produce power evaluation results within seconds. This is achieved by directly extracting our in-house evaluation results based on three public N6-methyladenosine datasets on GEO as the pilot data (Niu et al., 2013; Schwartz et al., 2014; Barbieri et al., 2017). Our package also comes with a vignette that provides thorough instructions and examples of its applications in differential analysis experimental design on N6-methyladenosine.

## Results

### Larger Sample Size Benefits DMR Detection

Under simulation settings outlined in Supplementary Materials S1, we next examine the relationship between sample size and power in DMR detection, given that determining sample size is a primary objective in our method. We adopt sample sizes of 2, 3, 5, 7, and 10 per group, and nominal FDR values of 0.05, 0.1, 0.15, and 0.2, both of which are common choices in MeRIP-seq experiments. The empirical results for major metrics are shown in Figure 2. Grouped by sample size and nominal FDR level, power, targeted power, FDC, and FDR averaged over 100 simulations are shown in Figure 2A-D. For a fixed sample size, metrics like power, FDC, and FDR diminish under lower FDR thresholds. This occurs as lower FDR values lead to greater stringency, which in turn reduces false positives. The power will drop, as expected, when using stringent FDRs. As sample size increases, these differences become smaller, particularly for statistical power (Figure 2A). Here, power remains consistently high across all FDR levels with 7 and 10 replicates per group. This highlights the benefit of using a larger sample size that helps detect DMRs with limited effect sizes, where a type II error would often occur when the sample size is small. Such trend is observed consistently when using different pilot data (Figure S1). At the same time, these results give researchers the knowledge to optimize the sample size based on their budgets. Using Figure 2A as an example, a power around 0.8 is achieved with 7 samples per group, and sample size of 7 is considered large in current MeRIP-seq studies. The benefit of expanding the sample size to 10 is marginal, but the associated costs could be significantly higher.

**Figure 2.**
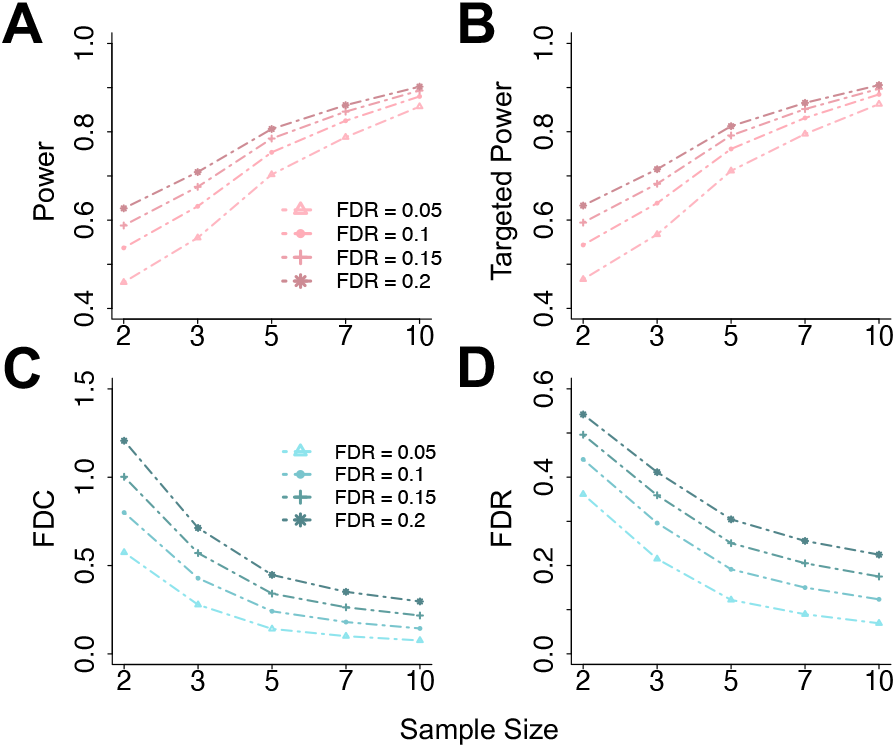
Statistical power evaluation metrics for DMR detection, at various sample sizes and FDR thresholds. (**A**) Power versus sample size, with each line presenting one FDR cutoff. (**B**)-(**D**) Similar to (**A**) but for other metrics: targeted power, FDC, and FDR. Targeted power and FDC are computed for DMRs with |*β*| ≥ 2. Each point on the line plots is an averaged value over N=100 simulations based on real MeRIP-seq data.

### Impact of Baseline Expression Values

It is useful for researchers to understand the effects of heterogeneity in baseline expression levels in DMR detection. In MeRIP-seq data, the basal expression level is represented by input control read counts, thus we stratify power metrics by input control ranges. Six strata are obtained based on following quantiles of mean input counts: stratum 1 (0%-10%), stratum 2 (10%-30%), stratum 3 (30%-50%), stratum 4 (50%-70%), stratum 5 (70%-90%), and stratum 6 (90%-100%). At a nominal FDR of 0.05, the average targeted power and FDC for the six strata are shown in Figure 3A, B. Overall, reduced targeted power is observed in the lower strata, a trend that is more evident when sample sizes are small. This is expected, as true differences in low-expressed regions are often obscured by noise, making DMRs harder to detect. A limited sample size further exacerbates this issue. This suggests the potential benefits of increasing the sequencing depth, particularly when biological replicates are limited and more samples are hard to obtain. Here, relatively low strata will enjoy the benefit of more drastic power improvement. Interestingly, higher FDCs are reported in the upper strata, suggesting that more false positives are detected per true discovery in these highly expressed regions. However, this trend diminishes with increasing sample size. Given that these metrics were computed across various simulation scenarios, we further explore the variability of the results, using visualizations within a specific stratum and sample size in Figure 3C-F. With elevated sample sizes, there is reduced variability in both targeted power and FDC (Figure 3C, D). This is not surprising since it is more likely to capture the true dispersion with more replicates, leading to more consistent power estimates. However, this trend is not observed across the strata at a fixed sample size, suggesting the benefit of increasing sample size over sequencing depth for more reliable inferences. Under a fixed sample size (Figure 3E, F), an upward trend is observed across the strata for both targeted power and FDC. This trend aligns with the observations in Figure 3A, B, though some variability is evident. A heatmap panel is also available to illustrate the stratified results (Figure S3).

**Figure 3.**
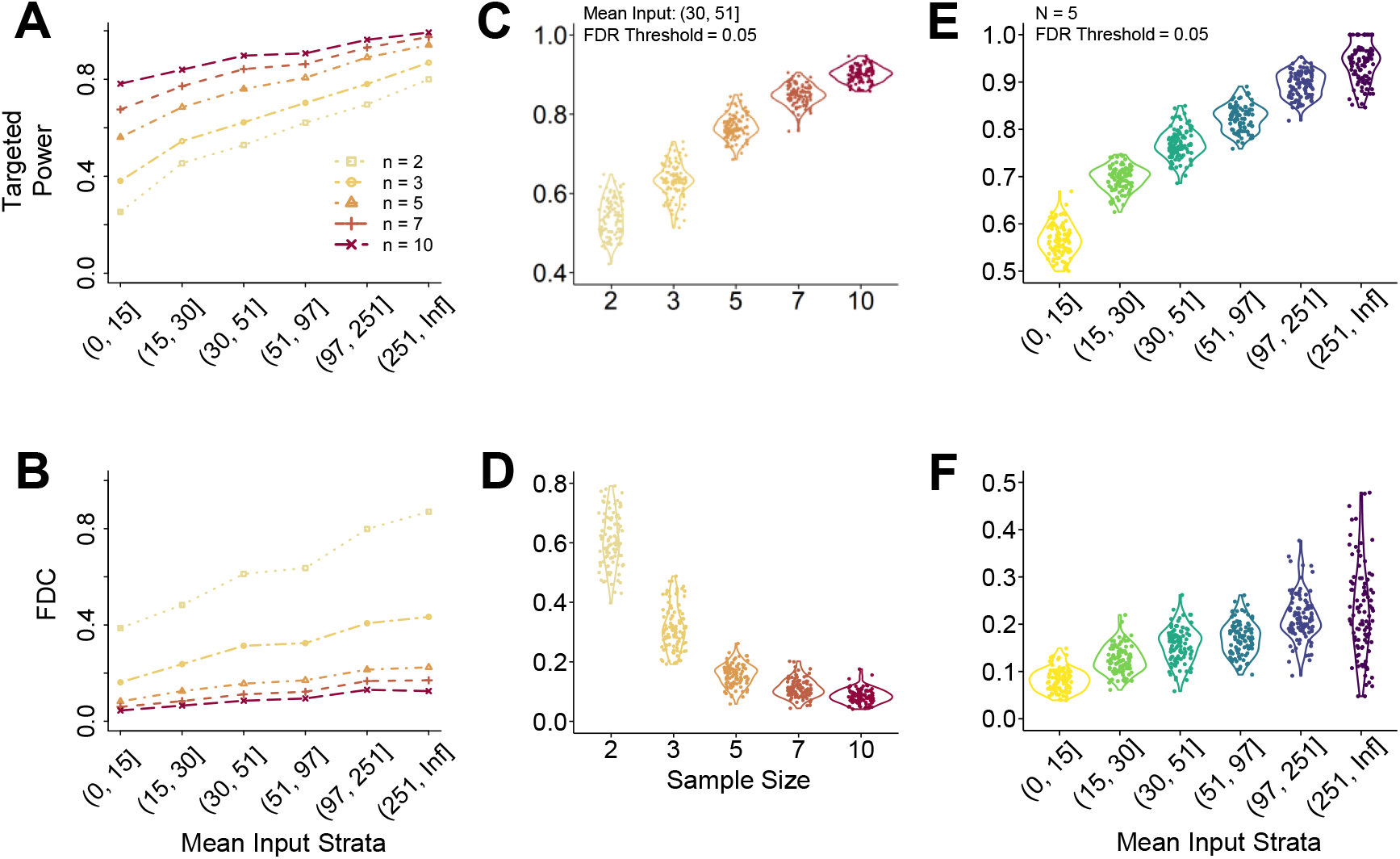
Targeted power and FDC stratified by mean input values for DMRs with |*β*| ≥ 2. Six strata are defined based on input count data quantiles: stratum 1 (0%, 10%), stratum 2 (10%, 30%), stratum 3 (30%, 50%), stratum 4 (50%, 70%), stratum 5 (70%, 90%), and stratum 6 (90%, 100%). A nominal FDR value of 0.05 is used to define significance. (**A**), (**B**) Mean targeted power and FDC along strata. Each line represents one sample size choice. (**C**), (**D**) Targeted power and FDC distributions in stratum 3, separated by sample size. (**E**), (**F**) Targeted power and FDC distributions with 5 replicates per group, stratified by mean input count values. N=100 simulations are conducted.

### Consistency Among Major DMR Calling Methods

It is worth noting that the targeted power and FDC presented in the Results section are computed for DMRs with odds ratios (OR) exceeding ∆ = 2, using TRESS. To evaluate the fluctuations of these two metrics across various effect sizes (OR), sample sizes and DMR detection methods, we also consider ∆ values of 1.5, 2, 4, 6, 8, and 10, for both TRESS and exomePeak2. In Figure 4, the targeted power and FDC are plotted against the sample size and are grouped by odds ratio thresholds. At all sample sizes, there is an increased targeted power (Figure 4A, C) and a higher FDC to identify DMRs with a larger odds ratio. Specifically, for FDC (Figure 4B, D), substantially higher values are observed among DMRs with exceptionally large odds ratios (∆ = 8, 10). This indicates that detecting DMRs with these large ORs might lead to a significant increase in false positives. These patterns hold true for both TRESS and exomePeak2. While both methods show improvements in targeted power with added replicates across all odds ratio thresholds, a discrepancy is noted for FDC that it tends to increase with larger sample sizes when using exomePeak2. This discrepancy, however, is not universally observed when applying our proposed framework to different pilot data sets (Figure S2). These findings highlight the importance of utilizing the users’ designated DMR detection methods during power calculation to ensure accurate estimations.

**Figure 4.**
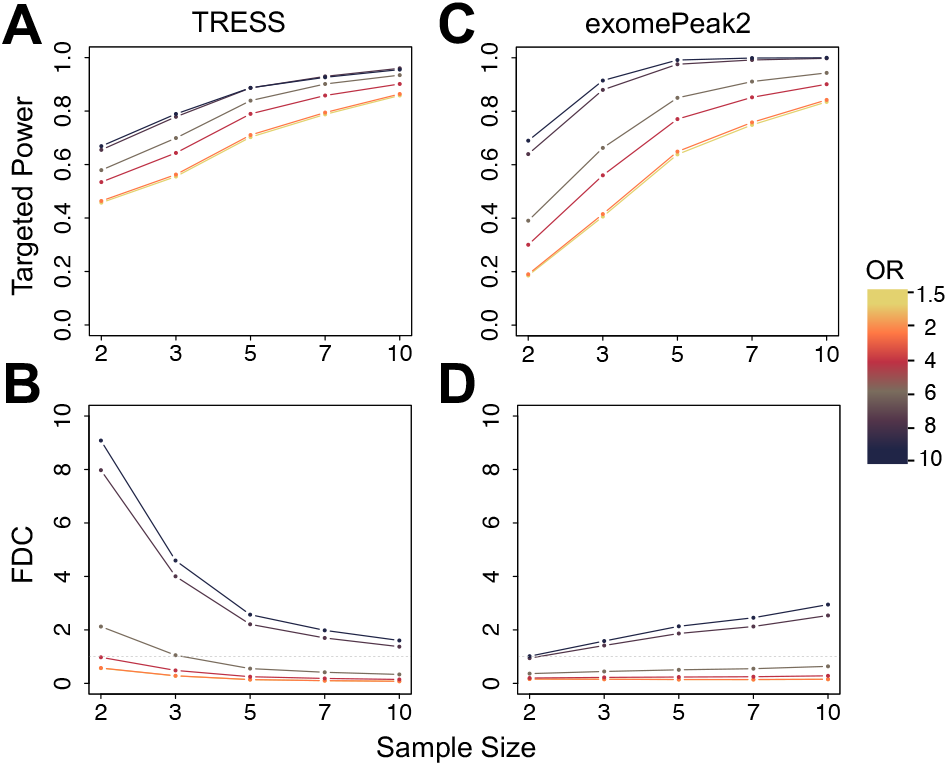
Comparing power evaluation results between major DMR detection methods TRESS (**A**)-(**B**) and exomePeak2 (**C**)-(**D**). Targeted power and FDC are shown at various Odds Ratios (OR, representing effect size) and sample sizes. A nominal FDR value of 0.05 is used to define significance. Points on the line plots are averaged over N=100 simulations.

### Impact of Sequencing Depth

As shown previously, sequencing depth is another critical factor in MeRIP-seq study design. Building upon our analysis of sequencing coverage strata, here we examine another aspect of sequencing depth by introducing a “depth factor”. This is a relative ratio to reflects the effect to enlarge or down-sample the sequencing coverage of the pilot data. As illustrated in Figure 5A, the targeted power rises with increased sequencing depth in all sample sizes. The incremental gain from increasing sequencing depth diminishes at high depths or large sample sizes, but benefits the small sample size the most. In Figure 5B for FDC, a similar pattern is observed as in the stratified analysis: FDCs increase with sequencing depth, but stabilize in scenarios with larger sample sizes. We also provide an integrated visualization in Figure 5C, presenting targeted power and FDC in the same panel, aiding users in understanding the tradeoffs between them. Researchers could consult similar figures, generated by ***magpie*** using their own pilot data, to select a customized increase in sequencing depth to achieve the desired power.

**Figure 5.**
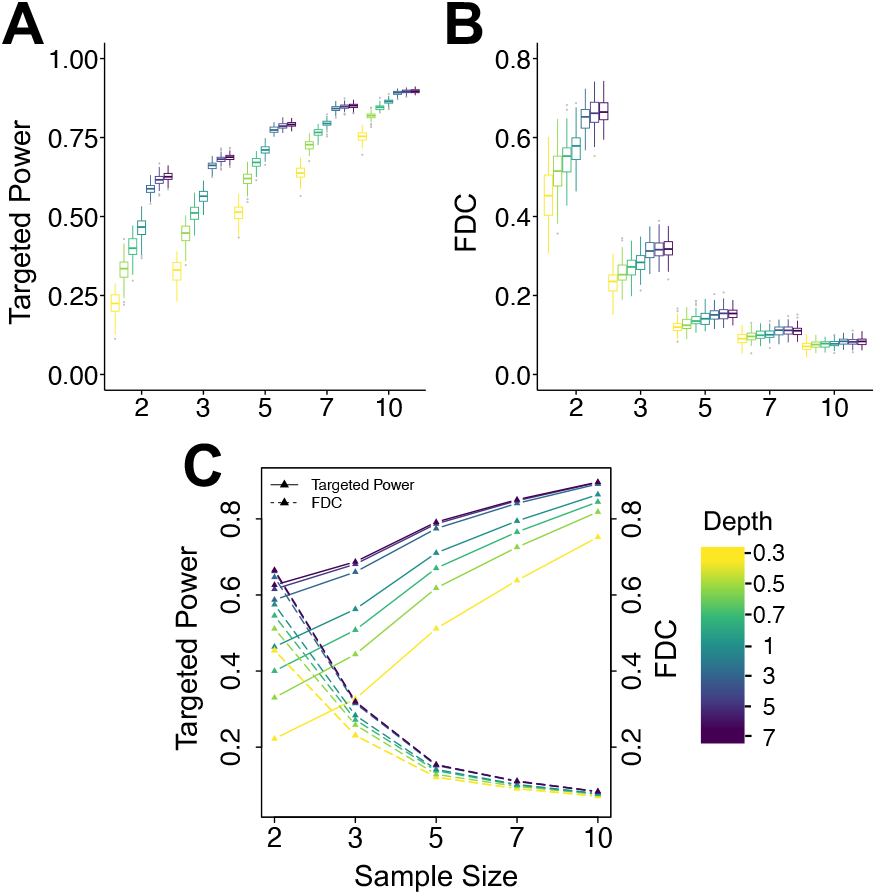
Sequencing depth affects targeted power and FDC for DMRs with |*β*| ≥ 2. The ‘depth factor’ is the relative ratio of the new dataset’s library size over that from the original dataset. It reflects the impact of enlarging or down-sampling the sequencing depth of pilot data. (**A**), (**B**) Targeted power and FDC under different sequencing depths, grouped by sample size. (**C**) Joint visualization of the mean targeted power and FDC, over various sequencing depths and sample sizes. N=100 simulations are conducted.

## Discussions

Sample size and power evaluation are pivotal and routine tasks in experimental design using sequencing data. Here we present the first tool to address the immediate needs of sample size calculation and power estimation for DMR detection in MeRIP-seq experiments. Traditionally, sample size calculation or power evaluation in hypothesis testings depends on inputs such as the effect size, variance from pilot studies, and the significance level. In contrast, for MeRIP-seq experiments with transcriptome-wide data, these scalar parameters must be considered as distributions. In addition, the distributions of sequencing depth and input control level can also significantly influence the statistical power, as we have shown in the results. We thus propose a statistically rigorous approach to address all these challenges, and draw information from pilot real data for simulation and empirical power evaluation.

We have a flexible simulation framework that allows switching models to mimic the real data well. In sequencing studies, data from varied tissues or cell types can exhibit unique expression and RNA methylation distributions across features (i.e. genes or regions). To address this, our tool allows users to provide pilot data analogous to their intended studies, serving as the basis for the estimated and adopted parameters in downstream simulations. To ensure that the simulated data accurately reflects actual data characteristics, ***magpie*** can adopt both negative-binomial and beta-binomial models and choose the one that aligns with the real data distributions best.

Both increased sequencing coverage and a larger sample size can significantly enhance the statistical power, as demonstrated in Results. Given that the total sequencing reads are often predetermined before experiments, researchers can benefit from our tool to optimize the balance between sequencing depth and sample size, to ensure the best possible experimental design in differential RNA methylation studies.

In our stratified analysis, significantly lower power is observed in regions of low input levels. This suggests the potential of refining the filtering strategy. While excluding low-expressed strata certainly means losing some true positives among these regions, it boosts the power to detect DMRs that are highly expressed, which are often of greater biological interest. Our proposed tool ***magpie*** can offer a foresight into the overall power gain, should the researchers want to weigh the tradeoffs before initiating their data analyses.

Our proposed approach captures real data characteristics, simulates data under various experimental settings, and produces common power evaluation metrices. This statistical framework has been implemented into a user-friendly R/Bioconductor package ***magpie***. The package allows users to save power evaluation results as an Excel file and visualize their relationship with aforementioned factors with line plots. Recognizing that users might not have their own pilot MeRIP-seq data, we also develop a “*quickPower*” function. This function can generate comprehensive power evaluation outputs in seconds, by retrieving pre-calculated results from three published studies. ***magpie*** is available at https://bioconductor.org/packages/magpie/.

## Acknowledgement

D.D., W.T., and H.F. were exclusively supported by intramural funding from Case Western Reserve University.

