## Supplementary Materials for "*magpie*: a power evaluation method for differential RNA methylation analysis in N6-methyladenosine sequencing"

Zhenxing Guo, Daoyu Duan, Wen Tang, Julia Zhu, William S. Bush, Liangliang Zhang, Xiaofeng Zhu, Fulai Jin and Hao Feng*

#### Table of Contents

**S1 Simulation Settings** **1**

**S2 Additional Results** **2**

#### S1 Simulation Settings

To demonstrate the power evaluation with our proposed framework, we based our simulations on samples from a GEO dataset (GSE114150) [1]. The data is obtained using the MeRIP-seq technique from eight major fetal tissues, revealing m6As related to tissue-specific activities. For DMR analysis, replicates from two experimental conditions are expected, therefore we incorporate liver and kidney samples to construct a 3 versus 3 design. Estimating parameters from this real data, we conduct 100 simulations on 10,000 candidate regions under different settings: replicates per group (2, 3, 5, 7, 10), FDR thresholds (0.05, 0.1, 0.15, 0.2), and factors of (0.3, 0.5, 0.7, 1, 3, 5, 7) proportional to the initial sequencing depth. Evaluation metrics are subsequently averaged for each scenario. We also employ four other pilot datasets (GSE120024, GSE46705, GSE47217, GSE48037) [2–5]for additional examinations.

### S2 Additional Results

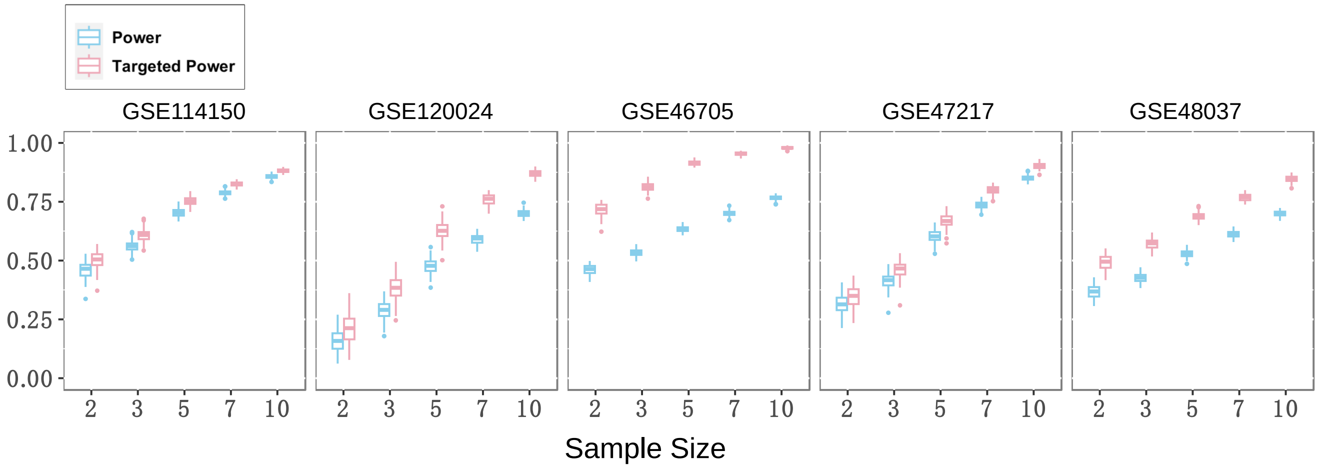


Figure S1: Boxplots comparing power and targeted power ($\Delta = 2$) across different sample sizes and pilot datasets. A nominal FDR value of 0.05 is used to define significance. N=100 simulations are conducted.

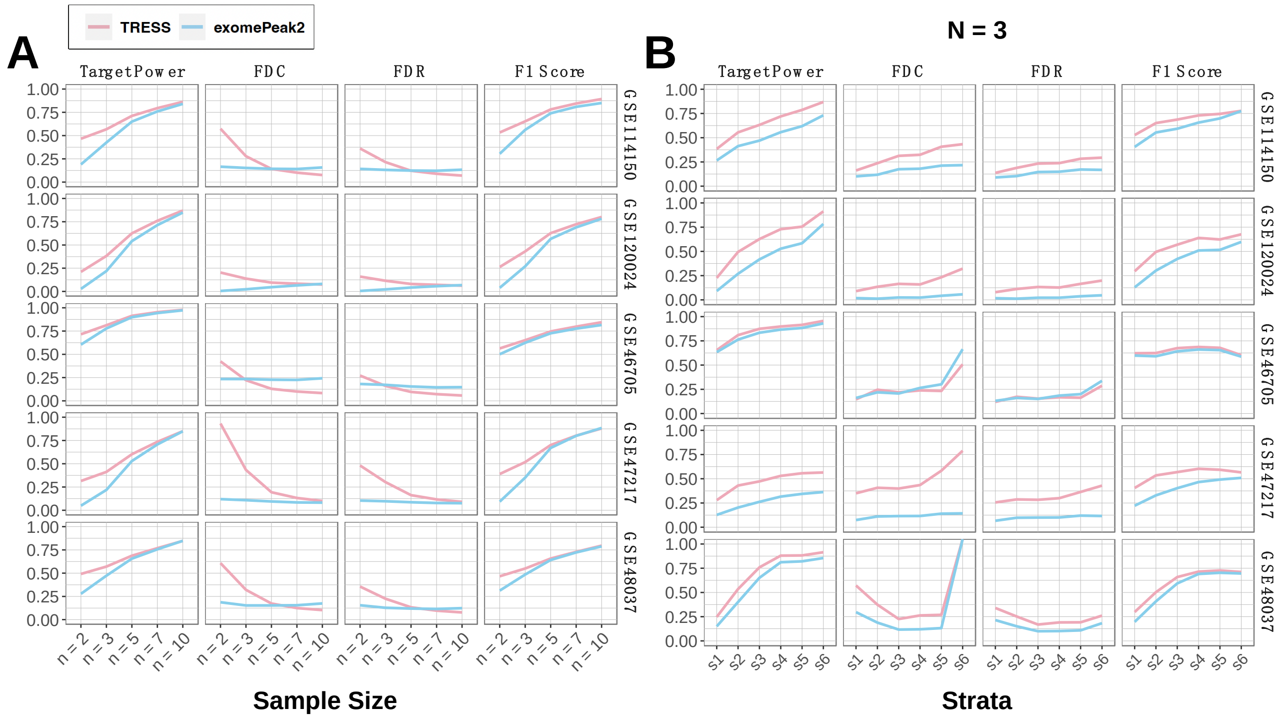


Figure S2: Comparing evaluation metrics across different pilot datasets. **A** Targeted power, FDC, FDR, and F score under sample size of 2, 3, 5, 7, 10 per group, across five pilot datasets. **B** The same metrics as **A**, but only for N = 3 and stratified by mean input counts. A nominal FDR value of 0.05 is used to define significance. N=100 simulations are conducted.

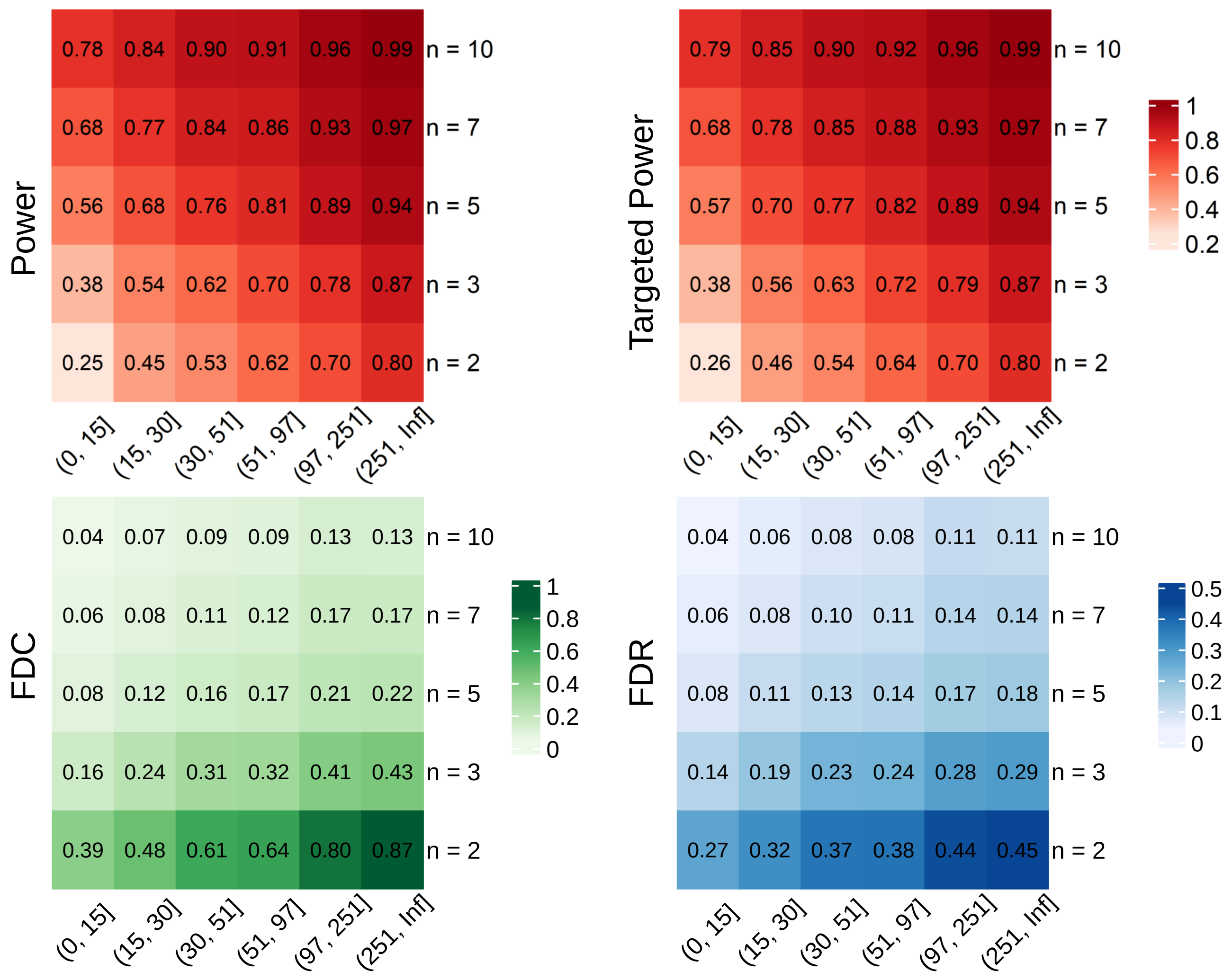


Figure S3: Heatmap showing power, targeted power, FDC and FDR stratified by mean input values. Six strata are defined based on input count data quantiles: stratum 1 (0%, 10%), stratum 2 (10%, 30%), stratum 3 (30%, 50%), stratum 4 (50%, 70%), stratum 5 (70%, 90%), and stratum 6 (90%, 100%). A nominal FDR value of 0.05 is used to define significance. N=100 simulations are conducted.
